# Invasive Earthworms Alter Forest Soil Microbiomes and Nitrogen Cycling

**DOI:** 10.1101/2021.03.07.433105

**Authors:** Jeonghwan Jang, Xianyi Xiong, Chang Liu, Kyungsoo Yoo, Satoshi Ishii

## Abstract

Northern hardwood forests in formerly glaciated areas had been free of earthworms until exotic European earthworms were introduced by human activities. The invasion of exotic earthworms is known to dramatically alter soil physical, geochemical, and biological properties, but its impacts on soil microbiomes are still unclear. Here we show that the invasive earthworms alter soil microbiomes and ecosystem functioning, especially for nitrogen cycling. We collected soil samples at different depths from three sites across an active earthworm invasion chronosequence in a hardwood forest in Minnesota, USA. We analyzed the structures and the functional potentials of the soil microbiomes by using amplicon sequencing, high-throughput nitrogen cycle gene quantification (NiCE chip), and shotgun metagenomics. Both the levels of earthworm invasion and soil depth influenced the microbiome structures. In the most recently and minimally invaded soils, *Nitrososphaera* and *Nitrospira* as well as the genes related to nitrification were more abundant than in the heavily invaded soils. By contrast, genes related to denitrification and nitrogen fixation were more abundant in the heavily invaded than the minimally invaded soils. Our results suggest that the N cycling in forest soils is mostly nitrification driven before earthworm invasion, whereas it becomes denitrification driven after earthworm invasion.

## Introduction

Earthworms are well-known ecosystem engineers that shape soil structure and drive nutrient dynamics in soil ecosystem [1]. They feed on litter and soil, burrow horizontally and vertically through soils, and release fecal materials to mix nutrients in soils, altering soil porosity, bulk density, water infiltration, gas emission, nutrient mineralization, and plant productivity [2].

Although earthworms are widely considered ubiquitous across the forest, grassland, agricultural, and garden ecosystems in the world, their global distribution is only beginning to be synthesized [3]. Glaciers and peri-glacial environments cleared out native earthworm populations from large areas in the northern USA and Canada as well as other Arctic areas in Eurasia during the last Ice Age [4]. Since then, most of these areas had remained earthworm-free until European earthworm species were introduced by human activities [5].

The earthworm invasion is now widely regarded as a force that substantially alters physical, geochemical, and biological properties of soils in northern hardwood forests [6, 7], and its ecosystem effects are believed to harm plant diversity [8] and be increasingly detrimental with ongoing changes in land uses and climates [9]. Invasive earthworms are known to reduce the litter layer (O horizon) while mixing organic matter with underlying minerals to create A horizon [10]. Presumably coupled with the loss of O horizon, invasion of European earthworms results in increased leaching of nitrates in the formerly glaciated deciduous forests [11]. Denitrification enzyme activity was also higher in the forest soils with earthworms than in those without earthworms [12].

A limited number of studies suggest that the invasion of earthworms could alter soil microbial communities. For example, Dempsey *et al.* [13] observed changes in soil microbial community composition based on the phospholipid fatty acid (PLFA) in a northern hardwood forest in New York, USA. Although PLFA analysis provides quantitative information, it cannot provide community compositions at low taxonomic (e.g., genus) levels. Hoeffner *et al.* [14] and de Menezes *et al.* [15] used terminal restriction fragment length polymorphism (T-RFLP) and high-throughput 16S rRNA gene amplicon sequencing, respectively, to analyze the effects of invasive earthworms on soil bacterial communities. However, these studies analyzed short-term impacts (e.g., 10-20 days [14] and 17 weeks [15]) by using soil mesocosms. Field investigation is essential to analyze the longer-term impacts of earthworm invasion in natural soil environments.

Previously, we studied the impacts of earthworm invasion on the soil physicochemical properties in a northern hardwood forest in Minnesota, USA [10, 16, 17], which built on decades-long research on ecological processes and effects of earthworm invasion [18–21]. At this site, a gradient of earthworm density was observed within a 190-m distance, which most likely reflects the history of earthworm invasion [17]. The invasion of earthworms has drastically changed the cycling of carbon, nitrogen, and other nutrients in soils [16, 17, 20]; however, it is unclear how it influenced soil microbial communities. Since microbes play crucial roles in soil C and N cycling in forest ecosystems [22], we hypothesize that the abundances of microbes important for C and N cycling have changed by the invasion of earthworms.

Consequently, the objectives of this study were to (1) elucidate the impacts of earthworm invasion on soil bacterial, archeal, and fungal communities at field conditions, (2) clarify the relationships between the levels of earthworm invasion, microbial communities, and soil physicochemical properties, and (3) analyze how earthworm invasion influenced the abundance of microbes/genes important for C and N cycles. To meet these objectives, we collected soil samples at different depths from three sites across an active earthworm invasion chronosequence in a hardwood forest in Minnesota. Our analyses, based on the amplicon sequencing, high-throughput nitrogen cycle gene quantification, and shotgun metagenomics, suggest that the structures and the functional potentials of the soil microbiomes altered by the invasion of earthworms. Microbial N cycling was most notably influenced. Our results suggest that the N cycling in forest soils is mostly nitrification driven before earthworm invasion, whereas it becomes denitrification driven after earthworm invasion.

## MATERIALS AND METHODS

### Soil sample collection

Soil samples were collected from a formerly glaciated northern hardwood forest in Minnesota, USA (Fig. S1). Earthworm biomass and species composition vary along a transect of ~200 m, while other environmental variables including climate, vegetation, geology, and topography are consistent within the transect as described in the supplementary information. We selected three sites along the transect: heavily invaded site (H), minimally invaded site (M), and the intermediate site (I). The minimally invaded site had the smallest earthworm biomass, dominated by epigeic earthworms (Fig. S2), which live and feed in a surface litter [23]. The heavily invaded site had the largest earthworm biomass, dominated by anecic and endogeic species. While endogeic earthworms burrow horizontally through soils and feed on decomposed matter and mineral soils, anecic earthworms live in deep vertical burrows and feed at a soil surface [23]. Although the total amount of earthworm biomass at the site I was similar to that at the site H, the population of anecic earthworms was small. Each ecological group (i.e., epigeic, endogeic, and anecic earthworms) differently influences soil ecosystems based on their feeding and moving behaviors [24].

After removing large undegraded leaves from the surface, three replicate soil cores (0-20 cm) were taken at each site by using a surface-disinfected soil probe. The soil core samples were divided into six segments (0-2 cm, 2-4 cm, 4-6 cm, 6-8 cm, 8-10 cm, and 10-20 cm by depth) and placed in Whirl-Pak bags. A total of 54 soil samples were collected (three sites × three soil cores × six segments). Samples were kept on ice immediately after collection, and frozen with dry ice within 2 h of collection. Soil physicochemical properties (Soil pH, bulk densities, and carbon, ammonium, nitrate, and nitrite contents) were measured in this study or obtained from previous literature as described in the supplementary information.

### DNA extraction, PCR, and amplicon sequencing

Total DNA was extracted from 0.25 g of each soil sample by using a DNeasy PowerSoil Kit (Qiagen) and QIAcube (Qiagen) according to the manufacturer’s instructions. From these DNA samples, the V4 region of the 16S rRNA gene and the fungal internal transcribes spacer 2 (ITS2) region between 5.8S and 23S rRNA gene were amplified and sequenced as described in the supplementary information.

The paired-end raw sequence reads were quality-filtered, trimmed, and assembled using NINJA-SHI7 [25]. The assembled sequences were clustered into operational taxonomic units (OTUs) at 97% sequence similarity by using NINJA-OPS [26]. Taxonomic assignments of the archaeal/bacterial and fungal OTUs were done using the Greengenes database version 97 [27] and UNITE [28] reference data sets, respectively. The resulting OTU tables with taxonomic information were used for statistical analyses (see below).

Based on the fungal OTU sequence data, fungal trophic modes and functional guilds were predicted by using the FUNGuild software [29]. Only results with the confidence scores of “Probable” and “Highly Probable” were used for statistical analyses.

### Nitrogen Cycle Evaluation (NiCE) chip

High-throughput microfluidic qPCR was used to quantify nitrogen cycle-associated genes (Nitrogen Cycle Evaluation [NiCE] chip) [30]. Several assays were newly added to the NiCE chip system to increase the target coverage. A total of 43 qPCR assays were included, targeting the genes associated with nitrification, denitrification, dissimilatory nitrate reduction to ammonium (DNRA), anaerobic ammonium oxidation (anammox), and nitrogen fixation (Table S1). Quantification was done using the standard curve method [31] as described in more detail in the supplementary information.

### Shotgun metagenomic sequencing

DNA extracted from surface soils (0-2 cm depth) were also used for shotgun metagenomics as described in the supplementary information. To identify N cycle-related genes, we mapped the high-quality metagenomic sequence reads against the NCycDB, a comprehensive nitrogen cycle protein sequence database [32], by using DIAMOND [33] with a minimum sequence identity of 70%, a minimum query coverage of 75%, and an E-value of <10^−5^. To identify fungal denitrification-associated genes, we used the reference sequences reported by Higgins et al. [34] for the read mapping. Description of the top-hit sequence (i.e., aligned with the lowest E-value) was extracted for each mapped read, grouped according to their potential functions, and used to create a read count table. The number of reads mapped to each of the sequence groups (i.e., gene functions) was normalized by the total number of high-quality sequence reads for each sample and used for statistical analyses.

High-quality metagenomic sequence reads were also mapped against the Greengenes and UNITE databases to identify bacterial/archaeal and fungal communities, respectively, by using bowtie2 which is implemented in the NINJA-OPS pipeline [26]. The resulting read count data were used for statistical analyses.

### Statistical analyses

Statistical significances in the quantitative data obtained in this study were tested with the Kruskal-Wallis rank-sum test by using R version 4.0.2 (https://www.r-project.org/). Microbial community structures were analyzed by using R with *vegan* [35], *phyloseq* [36], and *DESeq2* packages [37]. Principal coordinates analysis (PCoA) with Bray-Curtis distance matrices were used to visualize the dissimilarities in microbial communities among sites (i.e., the levels of earthworm invasion) and soil depth. Differences in microbial community structures were tested using permutational multivariate analysis of variance (PERMANOVA).

Canonical analysis of principal coordinates (CAP) was done using Bray-Curtis distance matrices to identify environmental variables associated with the patterns in microbial communities [38]. Environmental variables that were correlated with other variables at the Spearman’s ρ values of >0.80 were removed from the CAP analysis. Multicollinearity among the environmental variables was also identified by calculating variance inflation factors (VIF). Variables with a VIF of >10 were removed from the model in the CAP analysis. Furthermore, variables that were not significant (*p* >0.05) by PERMANOVA were removed from the model.

Taxa or genes that increased or decreased their relative abundance after earthworm invasion were identified by Spearman’s rank correlation analysis between gene abundance and earthworm biomass, which was the largest in the heavily invaded soil and the smallest in the minimally invaded soils (Fig. S2). In addition, differentially abundant taxa across samples were identified using *DESeq2* with α = 0.01.

### Nucleotide sequence accession numbers

The 16S rRNA gene and fungal ITS2 amplicon sequences as well as the shotgun metagenomics sequence reads were deposited to the GenBank database under the BioProject number PRJNA504043. The SRA accession numbers are available in Table S2 and S3.

## RESULTS

### Abundance of microbes in soils

The abundances of archaea/bacteria and fungi were estimated by qPCR targeting the 16S rRNA gene (Fig. 1A) and the ITS2 region (Fig. 1B), respectively. While abundances of archaea/bacteria and fungi were not significantly different by the levels of earthworm invasion (i.e., sites), they were significantly different by depth (*p* <0.01 by Kruskal-Wallis test). Both archaea/bacteria and fungi were most abundant in near-surface soils. Soil depth was negatively correlated with the copy numbers of the 16S rRNA gene (Spearman’s ρ = −0.77, *p* <0.01) and the ITS2 region (Spearman’s ρ = −0.60, *p* <0.05). Although differences in the abundance of archaea/bacteria between the surface soil (0-2 cm) and soils at 10-20 cm were similar by site, those between the surface soil and soils at 8-10 cm depth was the larger in the minimally invaded soil (site M) than heavily invaded soil (site H) (*p* <0.05 by Kruskal-Wallis test) (Table S4). The same trend was also seen for the differences in the fungal abundances between the surface soil (0-2 cm) and the soils at 8-10 cm (Table S4).

**Figure 1.**
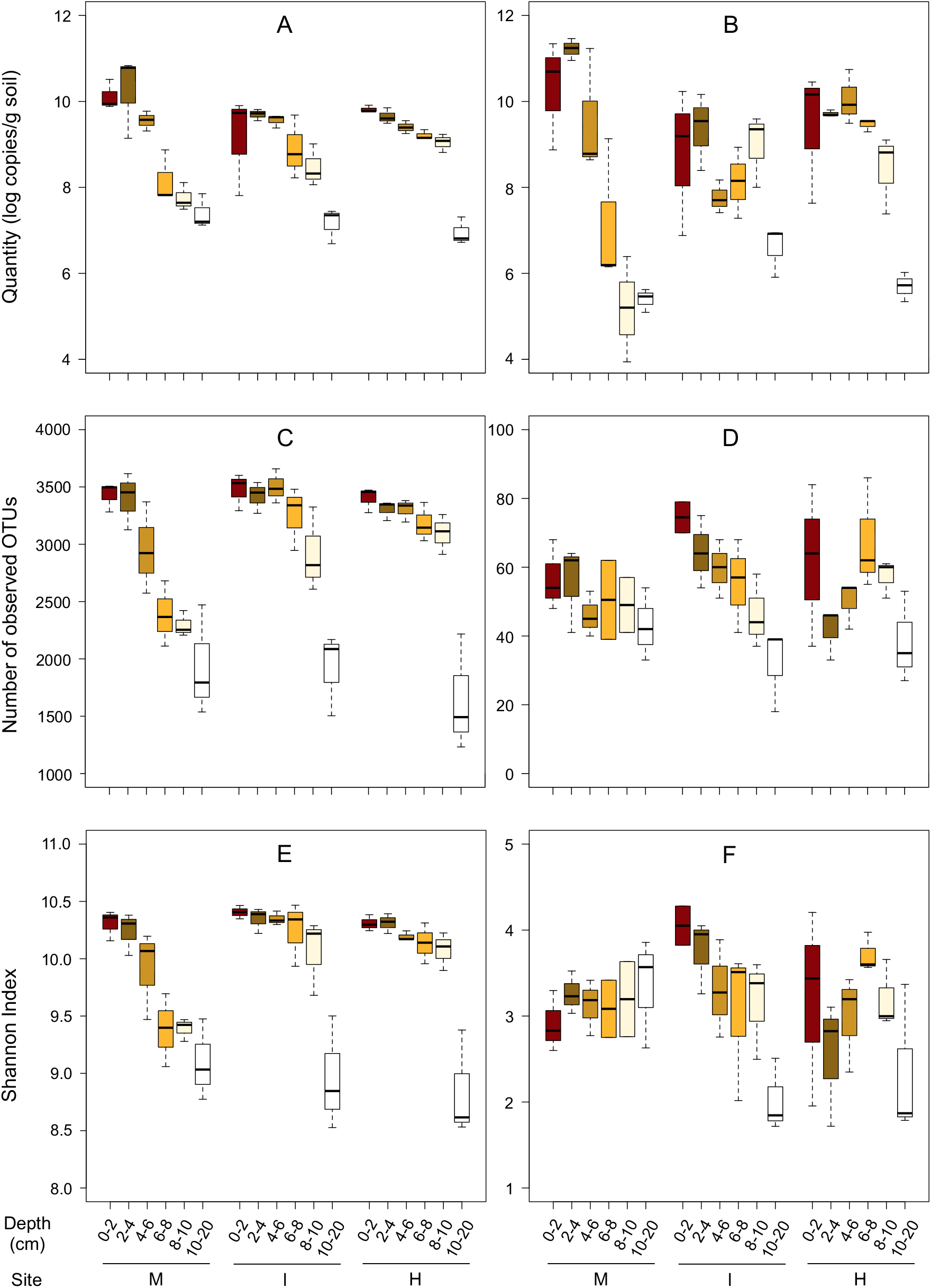
Abundance and diversities of microbes in the soil samples. Quantities of (A) 16S rRNA gene and (B) fungal ITS2 region measured by qPCR. Numbers of observed OTUs for (C) archaea/bacteria and (D) fungi. Shannon diversity index values calculated based on (E) the archaeal/bacterial 16S rRNA gene and (F) fungal ITS2 region sequences. Legend: M, the site with minimal invasion of earthworms; I, the site with an intermediate invasion of earthworms; and H, the site with a heavy invasion of earthworms.

### Alpha diversity measures

Soil archaeal/bacterial and fungal communities were analyzed by sequencing the 16S rRNA gene and ITS2 region, respectively. The number of sequences per sample ranged from 28 079 to 284 434 and from 967 to 53 610 for the 16S rRNA gene and ITS2 region, respectively (Table S2). Numbers of sequences were normalized at the smallest number of sequences by random subsampling for the diversity analyses. The subsampled sequences provided sufficient resolution of the microbial communities, as indicated by Good’s coverage ranging from 0.952 to 0.992.

Species richness was inferred by the numbers of observed OTUs for archaea/bacteria (Fig. 1C) and fungi (Fig. 1D). The numbers of archaeal/bacterial and fungal OTUs decreased by depth (*p* <0.01 and <0.05, respectively, by Kruskal-Wallis test), but not by the levels of earthworm invasion. Soil samples collected at 10-20 cm had the smallest number of OTUs for both archaea/bacteria and fungi, and the values were similar across sites. Differences in the numbers of observed OTUs for archaea/bacteria between the surface soil (0-2 cm) and the soils at 8-10 cm depth was larger in the site M than the sites H and I (Table S5).

Shannon diversity index values calculated based on the archaeal/bacterial 16S rRNA gene sequences were also significantly different by soil depth (*p* <0.01 by Kruskal-Wallis test) (Fig. 1E), whereas those calculated based on the fungal ITS2 region were not (Fig. 1F). Shannon index values for archaea/bacteria were the smallest in the soil samples collected at the 10-20 cm samples, and these values were similar across sites. Shannon diversity index values were not significantly different by the levels of earthworm invasion for both archaea/bacteria and fungi, although differences in the Shannon index values for archaea/bacteria between the surface soil (0-2 cm) and soils at 8-10 cm depth was greater in the site M than the sites H and I (Table S5).

### Patterns in soil microbial communities

Principal coordinates analysis (PCoA) showed the grouping of soil archaeal/bacterial communities by the levels of earthworm invasions (site M *vs*. site I and H) and soil depth (Fig. 2A). Community dissimilarities among different levels of earthworm invasions and soil depths were supported by PERMANOVA (*p* <0.01). The archaeal/bacterial communities in the near-surface soil (0-2 cm) and those in the deepest soil (10-20 cm) were most distantly plotted to each other. Distances between the archaeal/bacterial communities in the soils with minimum earthworm invasion (site M) and those in the soils with intermediate and heavy invasions (site I and H, respectively) were larger in the shallow soils (0-10 cm depths) than the deep soils (10-20 cm). Archaeal/bacterial communities in the deep soils overlapped each other, indicating that their community structures were similar to each other.

**Figure 2.**
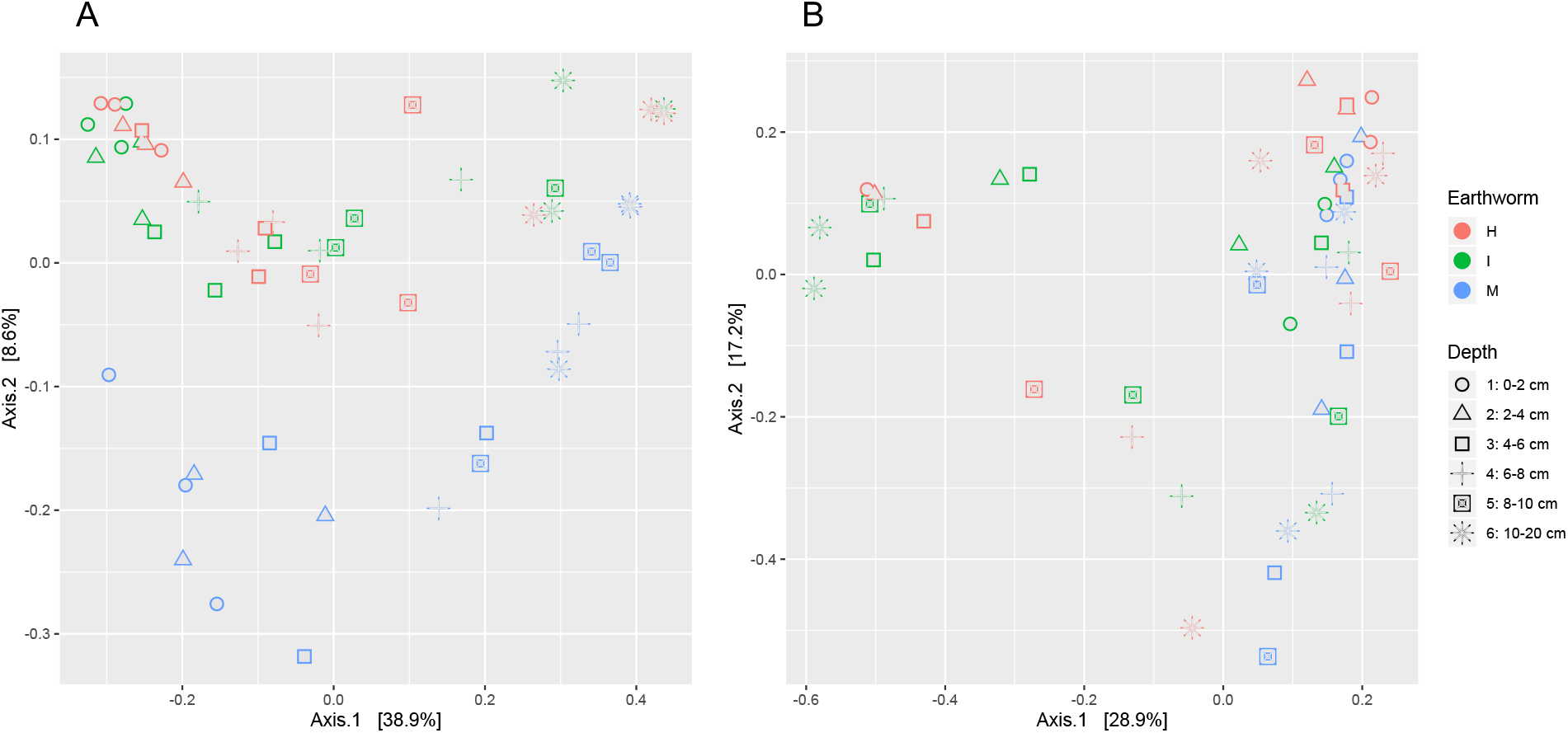
Principal coordinate analysis (PCoA) plots showing the Bray-Curtis dissimilarities among (A) archaeal/bacterial communities and (B) fungal communities. Each community is labeled with the site (M, the site with minimal invasion of earthworms; I, the site with an intermediate invasion of earthworms; and H, the site with a heavy invasion of earthworms) and soil depth (0-2, 2-4, 4-6, 6-8, 8-10, and 10-20 cm from the surface).

Fungal communities in the soils with different levels of earthworm invasion or different soil depths overlapped each other on the PCoA plot (Fig. 2B). However, fungal communities in the soils with minimum earthworm invasion were more closely related to each other than those in the soils with intermediate and heavy invasions, resulting in significant community dissimilarities by the levels of earthworm invasion (*p* <0.01 by PERMANOVA). In contrast, fungal communities were not different by soil depth (*p* =0.38 by PERMANOVA).

### Taxonomic composition

Major archaeal and bacterial phyla identified in this study include *Acidobacteria*, *Actinobacteria*, *Bacteroidetes*, *Crenarchaeota*, *Nitrospirae*, *Planctomycetes*, *Proteobacteria*, and *Verrucomicrobia* (Fig. 3A). Relative abundances of these phyla were similar among the soils with different levels of earthworm invasions, except for *Crenarchaeota* and *Nitrospirae.* Relative abundances of *Crenarchaeota* and *Nitrospirae* were the largest in the soil with minimal invasion of earthworms (site M) (*p* <0.01 by Kruskal-Wallis test). Similar results were obtained by the 16S rRNA gene amplicon sequencing and the shotgun metagenomics (Fig. S3).

**Figure 3.**
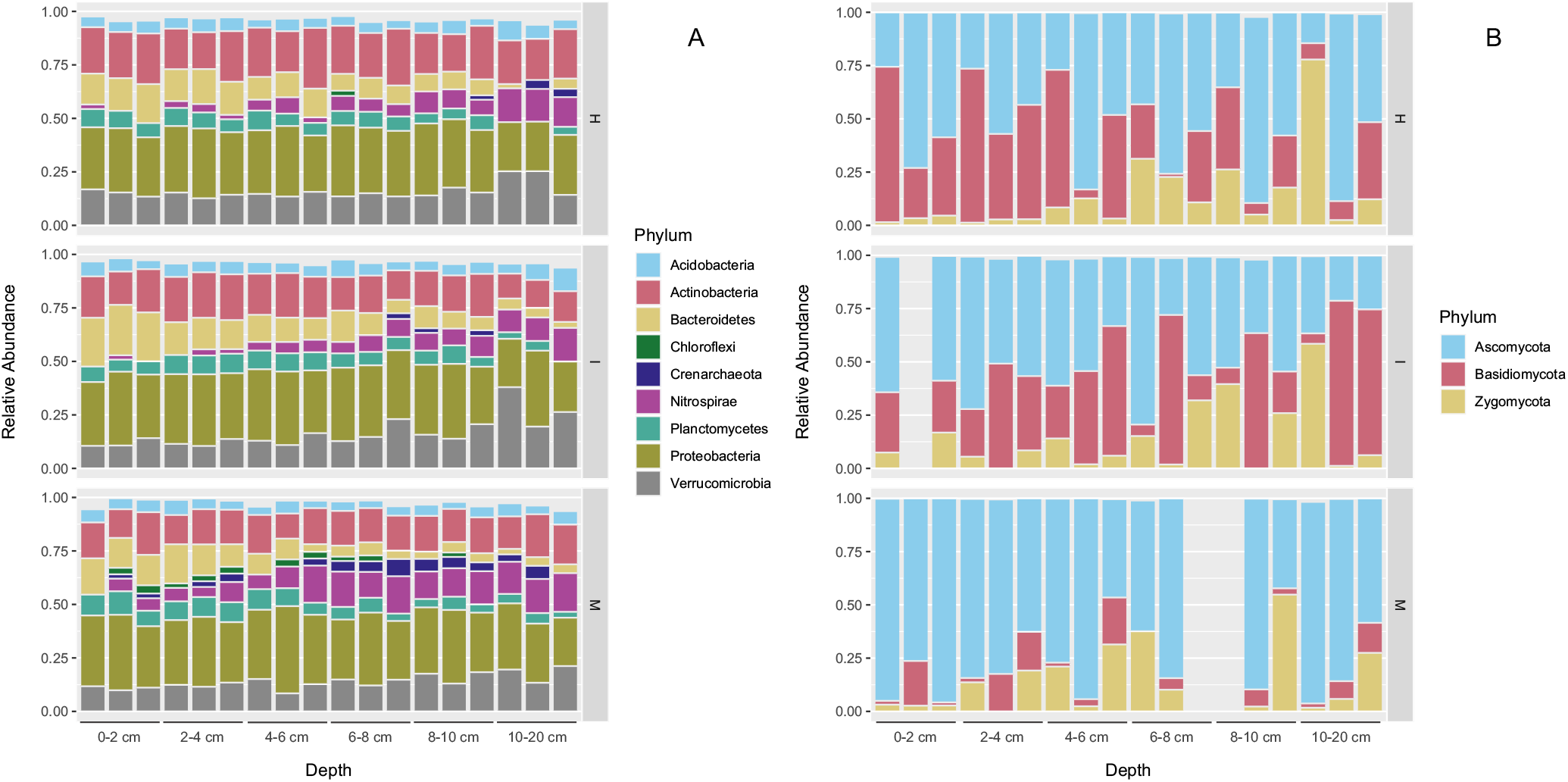
Relative abundance of (A) archaeal/bacterial phyla and (B) fungal phyla in the soil samples as assessed by the 16S rRNA gene and ITS2 sequencing analyses. Legend: M, the site with minimal invasion of earthworms; I, the site with an intermediate invasion of earthworms; and H, the site with a heavy invasion of earthworms.

*Ascomycota*, *Basidiomycota*, and *Zygomycota* were the major fungal phyla identified in this study (Fig. 3B). Relative abundances of *Basidiomycota* were larger in the soils with more earthworm invasions (site I and H) than those with minimal invasions earthworm invasions (*p* <0.01 by Kruskal-Wallis test).

### Microbial taxa responsive to earthworm invasion

To further identify the taxa that increased or decreased their relative abundance after the invasion of earthworms, Spearman’s rank correlation analysis was used (Table S6). Taxa that showed positive correlations with the levels of earthworm invasion included the genus *Mycobacterium* (ρ = 0.56, *p* <0.01) in the phylum *Actinobacteria*. Taxa that showed negative correlations with the levels of earthworm invasion included the genus *Nitrososphaera* (ρ = −0.60, *p* <0.01) in the phylum *Crenarchaeota*, the genus *Nitrospira* (ρ = −0.50, *p* <0.01) in the phylum *Nitrospirae*, and the fungal order *Helotiales* (ρ = −0.57, *p* <0.01) in the phylum *Ascomycota*.

### Quantities of the N cycle-associated genes

Some members of the genera *Nitrososphaera* and *Nitrospirae* play important roles in the N cycle, namely ammonia oxidation and nitrite oxidation, respectively. The decrease in their relative abundances in the earthworm-invaded soils could influence the overall N cycling in the soils. To clarify this, we used a high-throughput N-cycle gene quantification tool called the NiCE chip. With this tool, we could quantify almost all genes associated with the N cycle, including archaeal *amoA* and *nxrB* of *Nitrospira*. Of the 43 assays included in the NiCE chip, 18 assays showed quantitative results in >60% of the samples. These 18 assays targeted genes for nitrification (*amoA* and *nxrB*), denitrification (*napA*, *nirK*, *nirS*, *norB*, *nosZ*), and nitrogen fixation (*nifH*). In general, denitrification-related genes were more abundant than nitrification-related genes (Fig. 4A). Many of the denitrification- and nitrogen fixation-related genes were also more abundant in the soils with heavy earthworm invasions than the soils with minimal invasions (*p* <0.05 by Kruskal-Wallis test).

**Figure 4.**
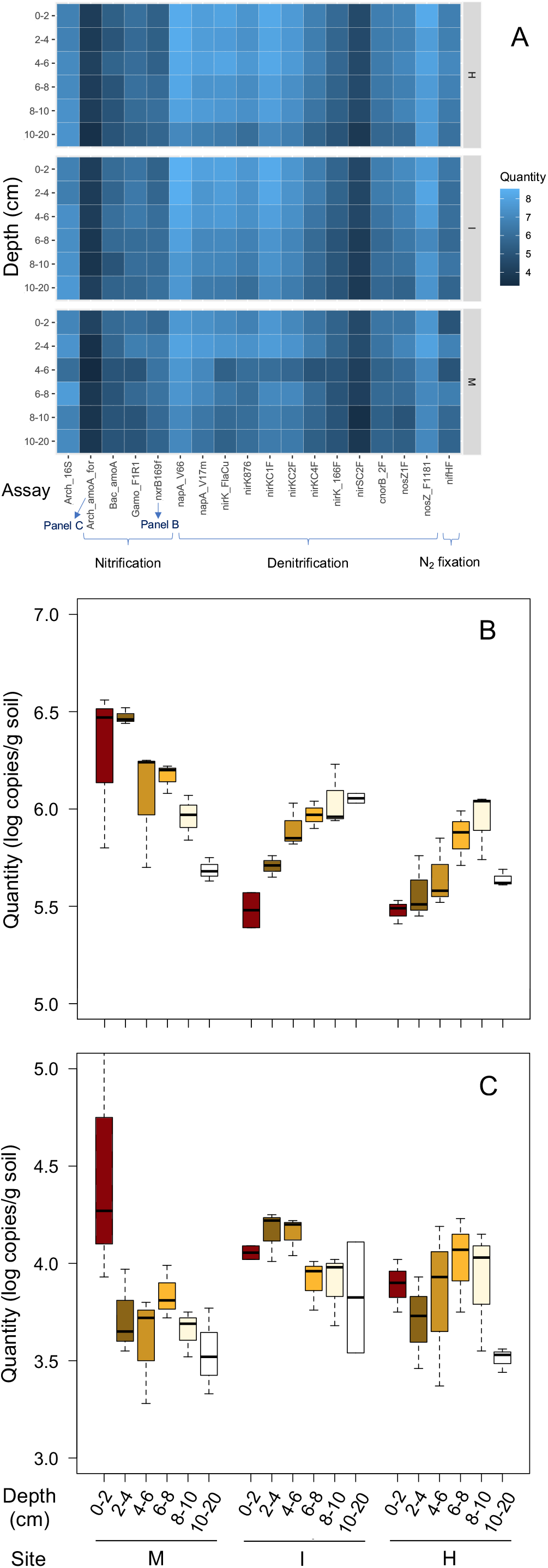
Quantities of N cycle-related genes in the soil samples. (A) Pseudo-heatmap showing the Nitrogen Cycle Evaluation (NiCE) chip results. Assays producing positives in >60% of samples are shown. (B) Quantities of nxrB of Nitrospira, based on the nxrB169f assay. (C) Quantities of archeal amoA based on the Arch_amoA_for assay. Legend: M, the site with minimal invasion of earthworms; I, the site with an intermediate invasion of earthworms; and H, the site with a heavy invasion of earthworms.

By contrast, the quantities of *nxrB* of *Nitrospira* (measured by the nxrB169f assay) were significantly greater in the soils with minimal invasion of earthworms than those heavily invaded by earthworms (*p* <0.01 by Kruskal-Wallis test) (Fig. 4B), in agreement with the changes in the relative abundance of *Nitrospira* measured by 16S amplicon sequencing. Interestingly, while quantities of *Nitrospira nxrB* decreased by depth in the samples with minimal invasion of earthworms (*p* <0.01 by Kruskal-Wallis test), those in the samples with heavy/intermediate invasions increased by depth (*p* <0.01 by Kruskal-Wallis test).

Quantities of archaeal *amoA* were not significantly different among the samples with different levels of earthworm invasion (Fig. 4C). In minimally invaded soils, archaeal *amoA* levels were significantly greater in the surface 0-2 cm soils than the soils collected at 10-20 cm depth (*p* <0.05 by Kruskal-Wallis and Mann’s *post hoc* test).

### Environmental variables associated with the patterns in microbial communities

Canonical analysis of principal coordinates (CAP) was used to clarify the relationship between environmental variables and the patterns in microbial communities. To select environmental variables for the CAP analysis, we first did a correlation analysis (Fig. S4). Most denitrification genes correlated with each other at Spearman’s ρ >0.8. Assay nirK_FlaCu was selected as the representative assay for denitrification because this assay produced quantitative values for all samples. Based on the correlation analysis, 13 variables were selected and used for the CAP analysis. After the VIF analysis and PERMANOVA, seven and five variables survived in the final CAP models for archaeal/bacterial and fungal communities, respectively (Fig. 5). Earthworm biomass, ammonium concentration, soil bulk density, and the quantities of denitrification and nitrogen fixation genes (nirK_FlaCu and nifHF assays, respectively) were commonly identified as the variables significantly associated with the patterns in both archaeal/bacterial and fungal communities (*p* <0.05 by PERMANOVA). For archaeal/bacterial communities, additional two variables, quantities of nitrification genes (Arch_amoA_for and nxrB169f) were also identified.

**Figure 5.**
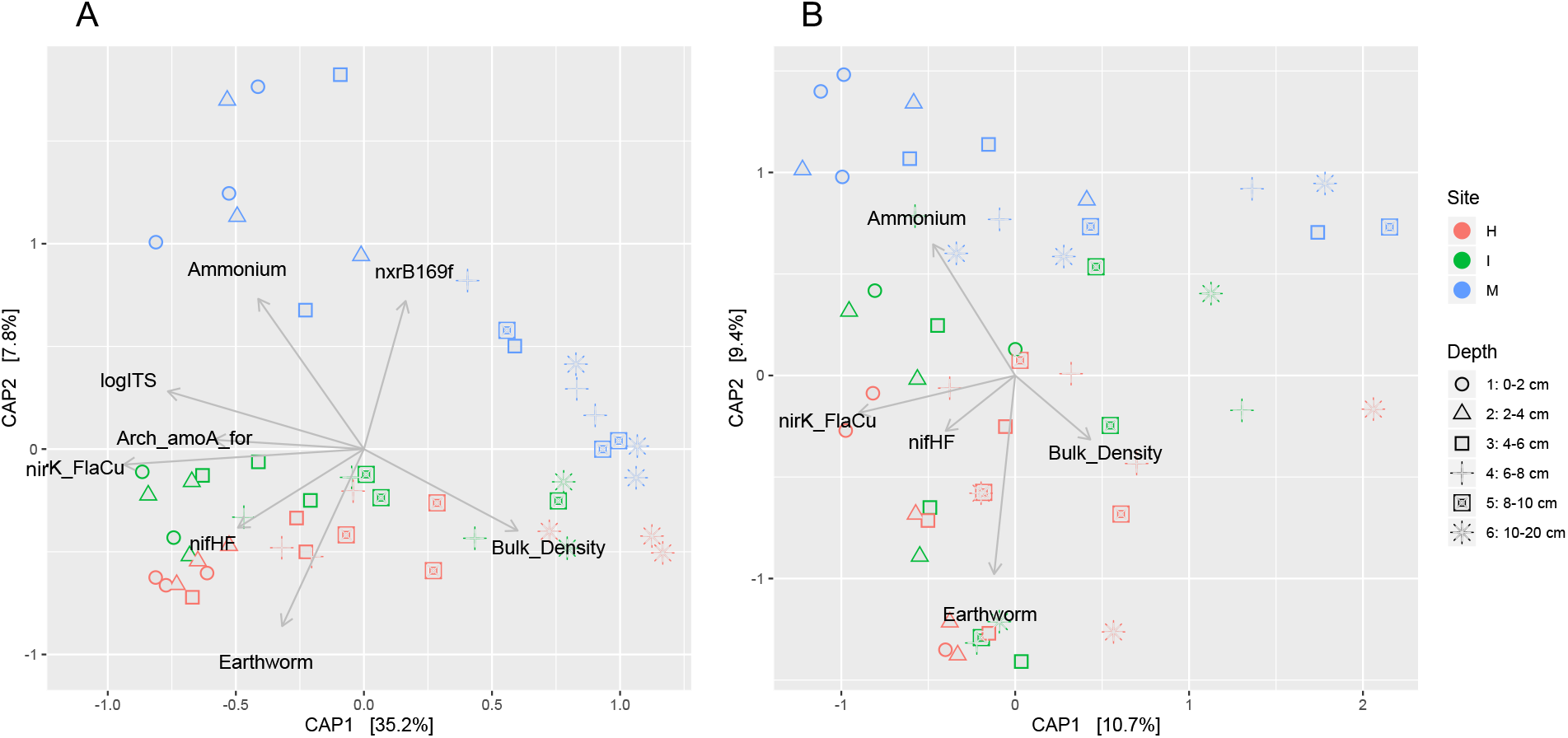
Canonical analysis of principal coordinates (CAP) plots showing the associations between environmental variables and the patterns in (A) archaeal/bacterial communities and (B) fungal communities. Microbial communities were analyzed using Bray-Curtis distance matrices. All environmental variables shown in the plots had significant effects (*p* <0.05) on the microbial community patterns based on PERMANOVA. Each community is labeled with the site (M, the site with minimal invasion of earthworms; I, the site with an intermediate invasion of earthworms; and H, the site with a heavy invasion of earthworms) and soil depth (0-2, 2-4, 4-6, 6-8, 8-10, and 10-20 cm from the surface).

### Functional potentials of the soil microbial communities

To further assess the functional potential of the soil microbial communities, we used the shotgun metagenomics approach. Surface soils (0-2 cm) were used for the metagenomic analysis because microbiomes in these soils were most significantly different between sites H, I, and M. Similar to the NiCE chip results, the relative abundances of the genes responsible for denitrification (*nar*, *nap*, *nor*, and *nos*) and nitrogen fixation (*nif*) were positively correlated with the levels of earthworm invasion (Spearman’s ρ >0.75, *p* <0.05) (Fig. 6). The relative abundances of bacterial/archaeal nitrite reductase genes (*nirK* and *nirS*) and fungal nitrite reductase gene (fungal *nirK*) were also positively correlated with the levels of earthworm invasion (Spearman’s ρ = 0.57 and 0.69, respectively), although they were not statistically significant (*p* = 0.143 and 0.057, respectively) (Table S7).

**Figure 6.**
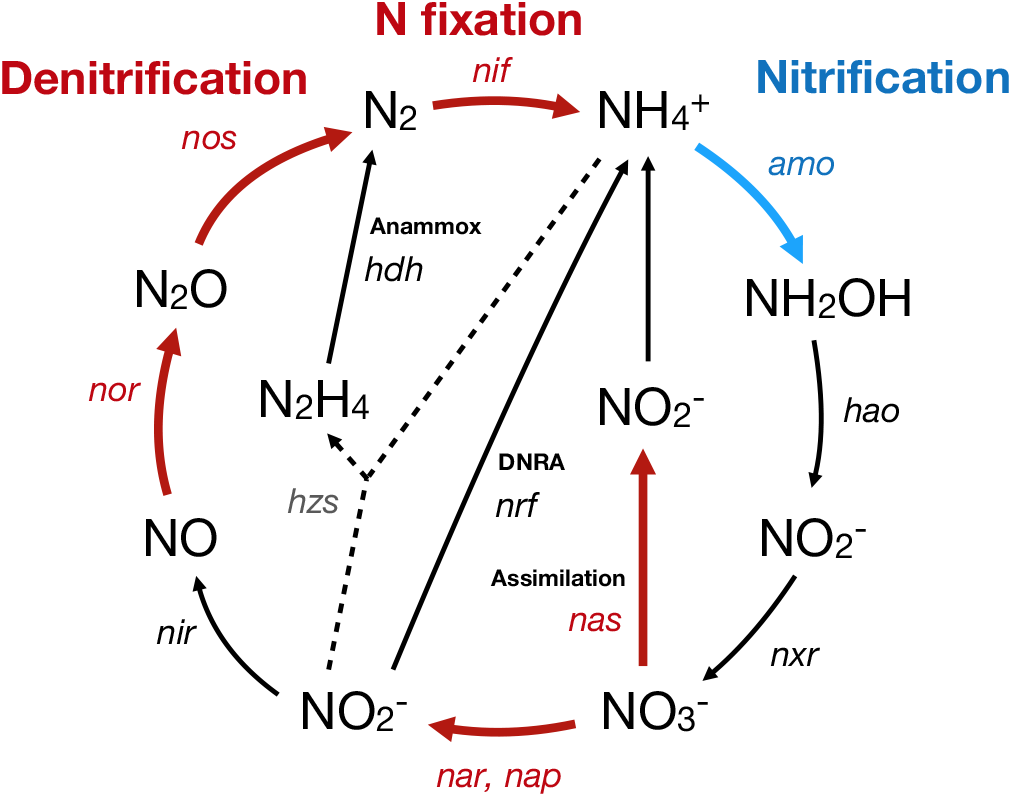
Nitrogen cycling in the forest soil influenced by earthworm invasion. Red and blue arrows indicate the genes that increased and decreased their relative abundances by earthworm invasion, respectively, based on Spearman’s correlation analysis of the shotgun metagenomics reads (*p* <0.05). Black solid arrows indicate the genes that did not change their abundance by earthworm invasion. Black dashed arrows indicate the genes that were not detected.

By contrast, the abundance of bacterial and archaeal ammonia monooxygenase genes for nitrification was negatively associated with the earthworm invasion levels (Spearman’s ρ < −0.80, *p* <0.05). The relative abundance of nitrite oxidoreductase gene (*nxr*) was also negatively yet insignificantly correlated with the levels of earthworm invasion (Spearman’s ρ = –0.57, *p* = 0.143). The relative abundance of glutamate dehydrogenase was positively correlated with the levels of earthworm invasion (Spearman’s ρ = 0.76, *p* <0.05), although other genes for nitrogen assimilation, such as glutamine synthetase or asparagine synthase were not (Table S7).

To assess the potential functions of the soil fungal communities, we used the FUNGuild approach. The relative abundance of symbiotrophs (e.g., ectomycorrhizae) was positively correlated with the levels of earthworm invasion (Spearman’s ρ >0.46, *p* <0.01); whereas those of saprotrophs and pathotrophs were negatively correlated with the levels of earthworm invasion (Spearman’s ρ < −0.41, *p* <0.01) (Table S8).

## Discussion

While the invasion of earthworms itself did not influence the abundances of archaea/bacteria and fungi as well as the α diversity measures, soil depths had significant impacts on the quantities and α diversities of soil microbiota. Effects of soil depths on microbial communities have been well documented in temperate forest soils [39, 40]. Impacts of soil depth were greater in the minimally invaded soil (site M) than more earthworm-invaded soils (sites H and I), as we found larger differences in the quantities and diversities of soil archaeal/bacterial populations between the surface (0-2 cm) and deep (8-10 cm) soils in minimally invaded soil than in heavily invaded soils. This is likely due to the mixing effects by soil-dwelling earthworms such as endogeic and anecic earthworms that were abundantly present in sites H and I. These earthworms vertically mix organic materials with minerals to form A horizon [16, 17]. Interestingly, differences in the quantities and α diversities of soil microbiota between the surface (0-2 cm) and the deepest (10-20 cm) soils were not significantly different by the level of earthworm invasion, suggesting that the impacts of earthworm invasion on soil microbial communities were minimal in the soils at 10-20 cm depth. This agrees with our field observations that A horizons rarely exceed 10 cm depth [17].

Both soil depth and the level of earthworm invasion had significant impacts on β diversity. The impact of earthworm invasion on β diversity of soil archaeal/bacterial communities was greater in shallow soils (0-10 cm) than in deep soils (10-20 cm). Indeed, archaeal/bacterial communities in the deep soils were similar to each other regardless of the levels of earthworm invasion. Although some anecic earthworms are known to dig deep burrows of up to 2 m [7], our results suggest that the invasion of earthworms had limited influence on the archaeal/bacterial community structures in the soils at 10-20 cm. This is also supported by the minimal presence of earthworm burrows below 10 cm as seen in the previous soil excavations to the depth of 1.5 m along the study transect including the site heavily infested with anecic *L. terrestris* [17]. The lack of deep earthworm activities may be due to the dry sandy loess layer and the dense clay-rich B horizon that underly the newly formed A horizon.

*Acidobacteria*, *Actinobacteria*, *Bacteroidetes*, *Planctomycetes*, *Proteobacteria*, and *Verrucomicrobia* occupied relatively large proportions of the archaeal and bacterial populations. Similar to this study, these phyla have been commonly detected in soils of temperate deciduous forests [41, 42]. Relatively large proportions of *Crenarchaeota* and *Nitrospirae* were also detected in this study, especially in the minimally invaded soils. These phyla are not frequently detected in other forest soils (e.g., [41, 42]), most of which presumably contained earthworms. *Nitrososphaera* spp. in the phylum *Crenarchaeota* and *Nitrospira* spp. in the phylum *Nitrospirae* were more abundant in the minimally invaded soils than the other soils. These microbes may play important roles in nitrification [43–46]. Interestingly, the vertical distribution of *Nitrospira* was different between the minimally invaded soils and more earthworm-invaded soils. *Nitrospira* spp. were more abundant near the surface of minimally invaded soils, probably because substrate for nitrification (ammonium and nitrite) were more available near the surface (O horizon). By contrast, *Nitrospira* spp. were more abundant in deeper soils of earthworm-invaded sites, probably because of the mixing soil microbes by earthworms and/or the leaching of nitrite downward.

While the relative abundance of *Nitrososphaera* spp. and *Nitrospira* spp. decreased by earthworm invasion, that of *Mycobacterium* (*Actinobacteria*) increased. Specific bacteria can be enriched in the earthworm guts, in which C, N, and other nutrients are more abundant than in soils [47, 48]. Indeed, *Mycobacterium* was frequently isolated from the guts of anecic earthworms (*L. terrestris*) [49] and was also identified as one of the bacterial taxa residing in earthworm gut walls of endogeic earthworms (*Aporrectodea* spp.) [24]. In addition, *Mycobacterium* was present at a significantly larger proportion in the guts of epi-endogeic earthworms (*L. rubellus*) than in soils [50]. These results collectively suggest that earthworms likely enriched *Mycobacterium* in their guts and contributed to the increased abundance of *Mycobacterium* in earthworm-invaded soils.

Metagenomic analysis revealed that archaeal *amoA* was more abundant in the minimally invaded soils than in the other soils, further supporting that archaea including *Nitrososphaera* spp. play ammonia oxidation in the minimally invaded soils. The qPCR analysis (i.e., NiCE chip), however, did not show a significant difference in the archaeal *amoA* abundance among the soils. Similarly, a discrepancy was also observed for the abundance of *nxrB* of *Nitrospira*: while NiCE chip results showed a significant difference in the abundance of *nxrB* of *Nitrospira* among the soils, no difference was detected by metagenomics. The discrepant results obtained by the two approaches may be due, in part, to (1) the large variations seen in the qPCR results, which was likely caused by the relatively low abundance of archaeal *amoA* and *Nitrospira nxrB* in the samples, (2) biases caused by the PCR primers (i.e., not all target genes are necessarily amplified by qPCR), or (3) the difference in the quantitative nature of the methods: while qPCR can provide absolute quantification (copies/g soil), metagenomics approach can provide only relative quantification (counts per million reads) in this study [51].

Both metagenomics and the NiCE chip analyses showed that many of the denitrification-related genes were more abundant in the earthworm-invaded soils than in the soils with minimal invasions. This may appear contradictory to the fact that the concentration of denitrification substrate (i.e., nitrate) was higher in the minimally invaded soil than the heavily invaded soils as seen in this study as well as Hale *et al.* [20]. However, it is probable that nitrate may have been consumed by denitrifiers in the earthworm-invaded soils, and therefore, became low compared to that in the minimally invaded soils. Similar to this study, greater denitrification activities were observed in earthworm-invaded forest soils than non-invaded soils [12, 52]. Higher denitrification activity was also reported in earthworm excreta (i.e., casts) than in the surrounding soils [53]. The larger abundance of denitrifying organisms in earthworm-invaded soils might be due, in part, to the enrichment of denitrifiers in earthworm intestines and/or the presence of more anoxic areas in soils. Earthworm intestines are known to promote denitrification, most likely due to the presence of anoxic area and readily assimilable carbon (i.e., mucus) [47, 48, 54, 55]. Earthworm-invaded soils could also have more anoxic micro-sites than the minimally invaded soils. Along our earthworm invasion chronosequence, both soil bulk density and the size and strength of soil aggregation are positively related with the level of earthworm invasion [16, 20], which is likely to offer more anoxic micro-sites that favor denitrification in the heavily invaded soils.

In addition to denitrification-related genes, the genes related to nitrogen fixation (*nif*) were also enriched in the earthworm-invaded soils in this study. The impact of earthworm invasion in nitrogen fixation is not well known; however, since total N content can decrease by earthworm-associated activities [56], the soil environment may become N-limited, which provides a selective advantage for N-fixing microbes.

Regarding the soil fungal communities, the relative abundance of *Basidiomycota* increased by earthworm invasion. This might be related to the increased abundance of symbiotrophs identified by the FUNGuild analysis because some fungi in *Basidiomycota* are known as being ectomycorrhizae and symbiotically associate with trees [57]. Ectomycorrhizae receive C mostly from their host plants instead of degrading complex organic matter in soils [57]. By contrast, the relative abundance of the order *Helotiales* (*Ascomycota*) decreased by the invasion of earthworms, which may be related to the decrease of the saprotrophs in the earthworm-rich soils. Most members of *Helotiales* live as soil saprophytes and degrade dead woods or other organic matter [58]. Similar to this study, Dempsey *et al.* [13] detected higher and lower levels of mycorrhizae and saprophytic fungi, respectively, in the earthworm-invaded soils than in the earthworm-free soils in a northern hardwood forest in New York, USA, by using the PLFA analysis. Collectively, these results suggest that soil fungal communities would shift from saprophytic to symbiotrophic communities by the invasion of earthworms, most likely due to the observed decrease in the organic C contents after earthworm invasions.

In conclusion, this study clearly shows that the invasion of earthworms alters soil microbial communities and ecosystem functioning. Earthworms mix soils, change soil physical structures, decrease the levels of C, N, and other nutrients in soils, secrete mucus, and enrich specific microbes in their guts, all of which can influence the microbial activities in soils. The most notable changes that earthworm invasion causes include the shift in the soil N cycling. Before the earthworm invasion, the N cycling in forest soils is mostly nitrification driven, for which AOA and *Nitrospira* play key roles. After earthworm invasion, the N cycling can become denitrification driven, which may cause N-limited conditions and the increased importance of the N fixing populations.

The invasion of earthworms is ongoing. They are actively spreading in formerly glaciated areas such as Alaska [59] and northern Scandinavia [60] in addition to Minnesota. To better understand the impacts of “global worming” on the ecology of soil systems, future research is necessary including the analysis of soil microbiomes at non-invaded and recently invaded soils on a global scale.

## Acknowledgements

We thank Marshal Landrum and Hao Wang for technical assistance, and Adrian Wackett for valuable comments. This research was supported by the MnDRIVE Initiative (to SI) and the CFANS Bridge & Development Grant (to KY and SI) of the University of Minnesota. This study was done, in part, by using the Minnesota Supercomputing Institute’s resources.

