## Supplementary Materials for "Invasive Earthworms Alter Forest Soil Microbiomes and Nitrogen Cycling"

This file contains supplementary methods (p. 2–7), eight tables (Table S1–S8, p. 8-19), four figures (Figure S1–S4, p. 20-23), and references (p. 24-26).

### Supplementary Methods

Soil sample collection

Soil samples were collected from a formerly glaciated northern hardwood forest near Leech Lake in north-central Minnesota, USA (Fig. S1). A detailed survey of soil physical properties and dendrochronology established that European earthworms arrived at the forest edge (recreational road) in the 1970s [1, 2] and have since invaded the forest for about 200 meters [3, 4]. It has been well established that environmental variables including climate, vegetation, geology, and topography are consistent within the transect as described in the supplementary materials.

Based on the earthworm biomass, species composition, and the time since the arrival of European earthworms, we selected three sites along the transect. The heavily invaded site (H) (47°16'0.5"N, 94°23'55.1"W, 390 m ASL) was the nearest from the presumed source of invasion (i.e., a road) and had the largest earthworm biomass (Fig. S2). The minimally invaded site (M) (47°15'59.9"N, 94°23'45.9"W, 411 m ASL) was located ~210 m from site H and had the smallest earthworm biomass. The intermediate site (I) (47°16'0.3"N, 94°23'48.4"W, 418 m ASL) was located between site H and site M (~150 m from site H). Although the total amount of earthworm biomass was similar between site H (5.40 ± 0.96 g/m$^2$) and site I (4.50 ± 0.68 g/m$^2$), species composition was different (Fig. S2) [3, 4]. While anecic earthworms (adult *Lumbricus terrestris*) were most abundant at site H, epi-anecic (juvenile *Lumbricus terrestris*) and endogeic (*Aporrectodea* spp. and *Octolasion* spp.) earthworms were more abundant at site M. Anecic earthworms were not observed at site M.

The soil type and plant vegetation were identical at these sites. The soil at these sites was mapped as the Warba soil series and classified as a fine-loamy, mixed, superactive, frigid Haplic Glossudalf (Soil Survey Staff 2014). However, organic matter contents were different most likely due to earthworm activities. At the H and I sites, O horizon was not observed; instead, A horizon was >5 cm [4]. At the M site, A horizon was thin (<5 cm), but the O horizon was >5 cm. The most dominant tree species at this forest was sugar maples (*Acer sacharuum*), but birch (*Betula papyrifera*, *Betula alleghaniensis*) and basswood (*Tilia americana*) also occurred.

Soil physicochemical parameters

Ammonium, nitrate, and nitrite contents in the soil profiles at each site were measured in this study. Ammonium ion was extracted from soil (2 g) by shaking with 30 ml of 2 M KCl for 30 min. Nitrate and nitrite ions were extracted from soil (2 g) by shaking with 30 mL 0.01 M $CaSO_4$ for 15 min. Ammonium, nitrate, and nitrite contents in the soil extracts were measured using a Lachat QuikChem 8500 Flow Injection Analyzer in the Soil Testing and Research Analytical Laboratory at the University of Minnesota. Soil pH, bulk densities, and carbon contents previously measured [3] were used in this study.

Amplicon sequencing

The V4 region of the 16S rRNA gene and the fungal internal transcribes spacer 2 (ITS2) region between 5.8S and 23S rRNA gene were amplified by using quantitative

PCR (qPCR) with Meta_V4_515F and Meta_V4_806R primer set [5] and 5.8SR_Nextera and ITS4_Nextera primer set [6], respectively. The qPCR mastermix was prepared using KAPA HiFidelity Hot Start Polymerase (Kapa Biosystems) as described previously [5]. The qPCR was done by using ABI 7900HT Real-Time PCR System (Applied Biosystem) with the following thermal conditions: 95°C for 5 min, followed by 35 cycles of 98°C for 20 sec, 55°C for 15 sec, and 72°C for 1 min, and one cycle of 72°C for 5 min. Melting curve analysis and agarose gel electrophoresis were done to confirm the correct amplification of the PCR products.

For sequencing library preparation, the V4 region of the 16S rRNA gene and the fungal ITS2 region were amplified as described above, except that the number of PCR cycles was decreased to 25 based on the qPCR results. This number of PCR cycles was set to reach the log-linear phase of the PCR kinetics to minimize PCR-dependent biases during amplification [7]. PCR products were purified and used to attach dual index tags and Illumina sequencing adaptors by PCR with Nextera kit as described previously in detail [5, 6]. The pooled and size-selected amplicons were diluted to 8 pM in Illumina's HT1 buffer. Paired-end sequencing reactions were done using a MiSeq platform (Illumina) with V3 chemistry (300-bp read length) at the University of Minnesota Genomics Center (UMGC).

<u>Nitrogen Cycle Evaluation (NiCE) chip</u>

High-throughput microfluidic qPCR was used to quantify nitrogen cycle-associated genes (Nitrogen Cycle Evaluation [NiCE] chip) [8]. Several assays were newly added to the NiCE chip system to increase the target coverage. A total of 43 qPCR assays were included, targeting the genes associated with nitrification, denitrification, dissimilatory nitrate reduction to ammonium (DNRA), anaerobic ammonium oxidation (anammox), and nitrogen fixation (Table S1).

The gBlock DNA fragments containing each of the target gene sequences were used as the standard DNA for the NiCE chip. When synthesis was difficult due to high GC contents, target gene fragments were PCR-amplified and cloned into a qCR 2.1 plasmid vector (Thermo Fisher). The plasmids were linearized and purified as described previously [9]. All of the standard DNA fragments were mixed and diluted to make a serial dilution of $5x10^6$, $5x10^5$, $5x10^4$, $5x10^3$, $5x10^2$, $5x10^1$, and $5x10^0$ copies/$\mu$l.

To increase the template DNA molecules, a preamplification (specific target amplification; STA) reaction was done prior to the NiCE chip run. The STA reaction is a multiplex PCR done with all primers with a small number of PCR cycles [9]. Both samples and standards were subjected to the STA reaction. The STA reaction mixture (8 $\mu$l) included 1x TaqMan PreAmp Master Mix (Thermo Fisher), 0.05 $\mu$M each primer, and 2 $\mu$l of template DNA. The STA reaction was done using a Veriti Thermal Cycler (Applied Biosystems) with the following condition: 95$^o$C for 10 min, followed by 14 cycles at 95$^o$C for 15 s and 60$^o$C for 4 min. To remove unreacted primers, 0.5 $\mu$l Exonuclease I (New England Biolabs) was added to each STA product and incubated at 37$^o$C for 30 min. The reaction was terminated by incubating the plate at 85$^o$C for 20 min. Reaction mixtures were then diluted to 40 $\mu$l with TE buffer (10 mM Tris-HCl and 0.1 mM EDTA [pH 8]).

The NiCE chip was run using a DynamicArray 96.96 GE chip and a BioMark HD System (Fluidigm). The sample premix (8 μl/sample) contained 1× SsoFast EvaGreen Supermix with low ROX (BioRad), 1X DNA Binding Dye (Fluidigm), and 3.6 μl of Exo 1-treated and diluted STA product. The assay premix (8 μl/assay) contained 1× Assay Loading Reagent (Fluidigm) and 5 μM each forward and reverse primers. The sample premix and assay premix were dispensed onto the DynamicArray 96.96 GE chip and mixed by using an IFC controller HX instrument (Fluidigm), according to the manufacturer's instructions. After mixing, the reaction mixtures contained 0.5 μM each forward and reverse primers. qPCR was performed under the following conditions: 50°C for 2 min, 70°C for 30 min, 25°C for 10 min, 95°C for 10 min, and 40 cycles of 96°C for 15 s, 50°C for 30 s, and 72°C for 1 min, followed by melting curve analysis from 50°C to 95°C at a rate of 1°C/3 s. ROX was used as a passive dye.

The quantification cycle ($C_q$) was determined for each assay by using Real-Time PCR Analysis Software version 3.0.2 (Fluidigm). The standard curves were generated by linear regression analysis of the $C_q$ values versus the known amounts of the standard DNA (log copies/μl) as described previously [9]. Quantities of target genes in the soil samples were calculated from the $C_q$ values (with 2 to 3 replicates) using the standard curves. Limit of quantification (LOQ) was defined as the lowest concentration of the standards that were reliably detected (2.7-3.7 log copies/μl). Assays were excluded from the downstream analysis when >40% of samples were below LOQ. When <40% of samples were below LOQ, these samples were given ½ LOQ values [10].

Shotgun metagenomic sequencing

DNA extracted from the surface soils (0-2 cm depth) at the three sites (H, I, and M) were also used for shotgun metagenomics. One of the replicates at the site I did not have enough amount of DNA, and therefore, was not used for metagenome analysis. As a result, eight samples were used for sequencing. Sequence libraries were created by using the Nextera XT kit (Illumina) with the mean library size of ~750 bp. Paired-end sequencing (150-bp read length) reaction was done using a NovaSeq platform (Illumina) at the UMGC. To ensure sequence quality for downstream analysis, Shi7 was used to pre-process raw metagenomic sequence reads [11]. Shi7 removed the 3′- and 5′-adaptors from sequence reads, and trimmed them at a minimum quality score of 25 and a minimum length of 50 bp.

High-quality metagenomic sequence reads were mapped against the Greengenes and UNITE databases to identify bacterial/archaeal and fungal communities, respectively, by using bowtie2 which is implemented in the NINJA-OPS pipeline [12]. The resulting read count data were used for statistical analyses as described in the Materials and Methods section. The identification of the N cycle-related genes is described in the Materials and Methods section

**Table S1.** Primers used for the high-throughput N cycle quantification (Nitrogen Cycle Evaluation [NiCE] chip)

| Target | Assay ID | Primer name | Sequence (5' --> 3') | Reference |
|---|---|---|---|---|
| Total Bacteria + Archaea | 16S | 515F | GTGCCAGCMGCCGCGGTAA | Herlemann *et al*. [13] |
| | | 806R | GGACTACHVGGGTWTCTAAT | |
| Total Archaea | Arch_16S | Archaea-F KO | CCCTAYGGGGYGCASCAGGC | Murakami *et al*. [14] |
| | | Archaea-R KO | GCYCYCCCGCCAATTCMTTTA | |
| AOB (Gamma-proteobacteria) | Gamo_F1R1 | Gamo172 F1 | GGBGACTGGGAYTTCTGG | Oshiki *et al*. [8] |
| | | Gamo172 F1_R1 | AAARCCCGAGAAGAAMGC | |
| | Gamo_F1R2 | Gamo172 F1 | GGBGACTGGGAYTTCTGG | |
| | | Gamo172 F1_R2 | AAAACCCGCAAAAAAGGC | |
| | Gamo_F2R1 | Gamo172 F2 | TGGGATTTCTGGATGGAC | |
| | | Gamo172 F2_R1 | TGATACGAACGCAGAGAA | |
| AOB (Beta-proteobacteria) | Bac_amoA | amoA_F1 | GGGGHTTYTACTGGTGGT | Rotthauwe *et al*. [15] Oshiki *et al*. [8] |
| | | amoA_2R | CCCCTCKGSAAAGCCTTCTTC | |
| AOB hao | hao | haoF4 | AYCTKCGCTCRATGGG | Schmid *et al*. [16] |
| | | haoR2 | GGTTGGTYTTCTGKCCGG | |
| AOA | Arch_amoAF | Arch-amoAF | STAATGGTCTGGCTTAGACG | Francis *et al*. [17] |
| | | Arch-amoAR | GCGGCCATCCATCTGTATGT | |
| | Arch_amoAFA | Arch-amoAFA | ACACCAGTTTGGYTACCWTCDGC | Beman *et al*. [18] |
| | | Arch-amoAR | GCGGCCATCCATCTGTATGT | |
| | Arch_amoAFB | Arch-amoAFB | CATCCRATGTGGATTCCATCDTG | |
| | | Arch-amoAR | GCGGCCATCCATCTGTATGT | |
| | Arch_amoA-for | Arch-amoA-for | CTGAYTGGGCYTGGACATC | Wuchter *et al*. [19] |
| | | Arch-amoA-rev | TTCTTCTTTGTTGCCCAGTA | |
| NOB (Nitrobacter) | nxrBF | NxrB 1F | ACGTGGAGACCAAGCCGGG | Vanparys *et al*. [20] |
| | | NxrB 1R | CCGTGCTGTTGAYCTCGTTGA | |
| | nxrB169f | nxrB169f | TACATGTGGTGGAACA | Pester *et al*. [21] |

| | | | | |
|---|---|---|---|---|
| NOB (Nitrospira) | | nxrB638r | CGGTTCTGGTCRATCA | |
| Comammox | comaA* | comaA-244F | TAYAAYTGGGTSAAYTA | Pjevac *et al.* [22] |
| | | comaA-659R | ARATCATSGTGCTRTG | |
| | comaB* | comaB-244F | TAYTTCTGGACRTTYTA | |
| | | comaB-659R | ARATCCARACDGTGTG | |
| Anammox | hzocl | hzocl1F1 | TGYAAGACYTGYCAYTGG | Li *et al.* [23] |
| | | hzocl1R2 | ACTCCAGATRTGCTGACC | |
| | hzsA | hzsA_1597F | WTYGGKTATCARTATGTAG | Harhangi *et al.* [24]; Oshiki *et al.* [8] |
| | | hzsA1857R | AAABGGYGAATCATARTGGC | |
| DNRA | nrfA | nrfAF2aw | CARTGYCAYGTBGARTA | Welsh *et al.* [25] |
| | | nrfAR1 | TWNGGCATRTGRCARTC | |
| Denitrification | narG_W9F* | W9F | MGNGGNTGYCCNMGNGGNGC | Gregory *et al.* [26] |
| | | T38R | ACRTCNGTYTGYTCNCCCCA | |
| | narG_1960f | narG1960f | AYGTSGGSCARGARAA | Philippot *et al.* [27] |
| | | narG2650r | TYTCRTACCABGTBGC | |
| | napA_V66 | V66 | TAYTTYYTNHSNAARATHATGTAYGG | Flanagan *et al.* [28] |
| | | V67 | DATNGGRTGCATYTCNGCCATRTT | |
| | napA_V17m | V17m | TGGACVATGGGYTTYAAYC | Bru *et al.* [29] |
| | | napA4r | ACYTCRCGHGCVGTRCCRCA | |
| | nirS_cd3aF | nirSCd3aF | AACGYSAAGGARACSGG | Kandeler *et al.* [30] |
| | | nirSR3cd | GASTTCGGRTGSGTCTTSAYGAA | |
| | nirSC1F* | nirSC1F | ATCGTCAACGTCAARGARACVGG | Wei *et al.* 2015 [31] |
| | | nirSC1R | TTCGGGTGCGTCTTSAGAASAG | |
| | nirSC2F | nirSC2F | TGGAGAACGCCGGNCARGTNTGG | |
| | | nirSC2R | GATGATGTCCACGGCNACRTANGG | |
| | nirSC3F* | nirSC3F | TTCGCCCTGAARGAYGGNGG | |
| | | nirSC3R | AGGTGCCCACGAANARNCCNCC | |

| | | | | |
|---|---|---|---|---|
| | nirK_FlaCu | FlaCu | ATCATGGTSCTGCCGCG | Throbäck *et al.* [32] |
| | | R3Cu | GCCTCGATCAGRTTGTGGTT | |
| | nirK876 | nirK876 | ATYGGCGGVAYGGCGA | Henry *et al.* [33] |
| | | nirK1040 | GCCTCGATCAGRTTRTGGTT | |
| | nirKC1F | nirKC1F | ATGGCGCCATCatggtnytncc | Wei *et al.* [31] |
| | | nirKC1R | TCGAAGGCCTCGatnarrttrtg | |
| | nirKC2F | nirKC2F | TGCACATCGCCAACggnatgtwygg | |
| | | nirKC2R | GGCGCGGAAGATGshrtgrtcnac | |
| | nirKC4F | nirKC4F | TACGGTGTGATCatcrtsgatcc | |
| | | nirKC4R | GCATCACGCATGgaatgatysac | |
| | norB2 | norB2 | GACAARHWVTAYTGGTGGT | Casciotti and Ward [34] |
| | | norB6 | TGCAKSARRCCCCABACBCC | |
| | cnorB-2F | cnorB-2F | GACAAGNNNTACTGGTGGT | Braker and Tiedje [35] |
| | | cnorB-6R | GAANCCCCANACNCCNGC | |
| | qnorB2F-5R | qnorB2F | GGNCAYCARGGNTAYGA | |
| | | qnorB5R | ACCCANAGRTGNACNACCCACCA | |
| | qnorB2F-7R* | qnorB2F | GGNCAYCARGGNTAYGA | |
| | | qnorB7R | GGNGGRTTDATCADGAANCC | |
| | nosZ1F | nosZ1F | WCSYTGTTCMTCGACAGCCAG | Henry *et al.* [36] |
| | | nosZ1R | ATGTCGATCARCTGVKCRTTYTC | |
| | nosZ-F-1181 | nosZ-F-1181 | CGCTGTTCITCGACAGYCAG | Rich *et al.* [37] |
| | | nosZ-R-1880 | ATGTGCAKIGCRTGGCAGAA | |
| | nosZ-II-F* | nosZ-II-F | CTIGGICCIYTKCAYAC | Jones *et al.* [38] |
| | | nosZ-II-R | GCIGARCARAAITCBGTRC | |
| | nosZ912F | NosZ912F | CGTCCCCGGCCTCGTGTA | Sanford *et al.* [39] |
| | | NosZ1853R | GAGCAGAAGTTCGTGCAGTAGTAGGG | |
| Nitrifier denitrification | nirK_166F | nirK_166F | GTWCCSGGTCCGGTYGTRCG | Cantera and Stein [40] |
| | | nirK_665R | TCRTTGGGWCCRGCRTTGAC | |

| | | | | |
|---|---|---|---|---|
| Fungal denitrification | nirKfF* | nirKfF | TACGGGCTCATGTAYGTNSARCC | Wei *et al*. [41] |
| | | nirKfR | AGGAATCCCACASCNCCYTTNTC | |
| N fixation | nifHF | nifHF | AAAGGYGGWATCGGYAARTCCACCAC | Rösch *et al*. [42] |
| | | nifHR | TTGTTSGCSGCRTACATSGCCATCAT | |
| | nifH_IGK3* | IGK3 | GCIWTHTAYGGIAARGGIGGIATHGGIAA | Ando *et al*. [43] |
| | | DVV | ATIGCRAAICCICCRCAIACIACRTC | |

* These assays did not produce amplicons in the NiCE chip format.

**Table S2.** Number of sequences obtained by the 16S rRNA gene and fungal ITS amplicon sequencing analyses (GenBank BioProject number PRJNA504043). Short read archive (SRA) accession numbers are also shown for each sample.

| Sample ID | Site | Depth | 16S rRNA gene | | Fungal ITS | |
|---|---|---|---|---|---|---|
| | | | Accession | No. of Seqs | Accession | No. of Seqs |
| MN.1.01 | M | 0-2 cm | SRR8166100 | 59195 | SRR8166165 | 17854 |
| MN.1.02 | M | 0-2 cm | SRR8166101 | 83388 | SRR8166164 | 12515 |
| MN.1.03 | M | 0-2 cm | SRR8166098 | 112871 | SRR8166163 | 14154 |
| MN.1.04 | M | 2-4 cm | SRR8166099 | 46550 | SRR8166162 | 11952 |
| MN.1.05 | M | 2-4 cm | SRR8166096 | 70808 | SRR8166161 | 15337 |
| MN.1.06 | M | 2-4 cm | SRR8166123 | 101751 | SRR8166160 | 16912 |
| MN.1.07 | M | 4-6 cm | SRR8166094 | 35482 | SRR8166159 | 18785 |
| MN.1.08 | M | 4-6 cm | SRR8166095 | 106899 | SRR8166158 | 8867 |
| MN.1.09 | M | 4-6 cm | SRR8166092 | 33588 | SRR8166157 | 14554 |
| MN.1.10 | M | 6-8 cm | SRR8166093 | 147837 | SRR8166156 | 10924 |
| MN.1.11 | M | 6-8 cm | SRR8166113 | 189231 | SRR8166199 | 10275 |
| MN.1.12 | M | 6-8 cm | SRR8166114 | 66833 | SRR8166198 | Not used[a] |
| MN.1.13 | M | 8-10 cm | SRR8166111 | 67747 | SRR8166201 | Not used[a] |
| MN.1.14 | M | 8-10 cm | SRR8166112 | 120702 | SRR8166200 | 5565 |
| MN.1.15 | M | 8-10 cm | SRR8166109 | 192389 | SRR8166195 | 8032 |
| MN.1.16 | M | 10-20 cm | SRR8166110 | 86473 | SRR8166194 | 3975 |
| MN.1.17 | M | 10-20 cm | SRR8166107 | 66592 | SRR8166197 | 9718 |
| MN.1.18 | M | 10-20 cm | SRR8166108 | 141055 | SRR8166196 | 1941 |
| MN.2.01 | I | 0-2 cm | SRR8166121 | 150716 | SRR8166207 | 11426 |
| MN.2.02 | I | 0-2 cm | SRR8166122 | 67707 | SRR8166206 | Not used[a] |
| MN.2.03 | I | 0-2 cm | SRR8166144 | 145194 | SRR8166179 | 10835 |
| MN.2.04 | I | 2-4 cm | SRR8166143 | 211957 | SRR8166180 | 14747 |
| MN.2.05 | I | 2-4 cm | SRR8166142 | 201105 | SRR8166181 | 20361 |
| MN.2.06 | I | 2-4 cm | SRR8166141 | 63255 | SRR8166182 | 25122 |
| MN.2.07 | I | 4-6 cm | SRR8166140 | 53775 | SRR8166183 | 20793 |
| MN.2.08 | I | 4-6 cm | SRR8166139 | 284434 | SRR8166184 | 21612 |
| MN.2.09 | I | 4-6 cm | SRR8166138 | 269897 | SRR8166185 | 52874 |
| MN.2.10 | I | 6-8 cm | SRR8166137 | 30063 | SRR8166186 | 25846 |
| MN.2.11 | I | 6-8 cm | SRR8166136 | 115118 | SRR8166177 | 25267 |
| MN.2.12 | I | 6-8 cm | SRR8166135 | 115106 | SRR8166178 | 13266 |
| MN.2.13 | I | 8-10 cm | SRR8166115 | 140831 | SRR8166209 | 11511 |
| MN.2.14 | I | 8-10 cm | SRR8166116 | 89291 | SRR8166208 | 39551 |
| MN.2.15 | I | 8-10 cm | SRR8166117 | 119878 | SRR8166166 | 36142 |
| MN.2.16 | I | 10-20 cm | SRR8166118 | 246344 | SRR8166193 | 13000 |

| | | | | | | |
|---|---|---|---|---|---|---|
| MN.2.17 | I | 10-20 cm | SRR8166119 | 170175 | SRR8166205 | 53610 |
| MN.2.18 | I | 10-20 cm | SRR8166120 | 28079 | SRR8166204 | 36266 |
| MN.3.01 | H | 0-2 cm | SRR8166126 | 233764 | SRR8166203 | 23668 |
| MN.3.02 | H | 0-2 cm | SRR8166097 | 131474 | SRR8166202 | 5798 |
| MN.3.03 | H | 0-2 cm | SRR8166102 | 158441 | SRR8166188 | 18184 |
| MN.3.04 | H | 2-4 cm | SRR8166105 | 156615 | SRR8166187 | 37118 |
| MN.3.05 | H | 2-4 cm | SRR8166128 | 181420 | SRR8166171 | 28942 |
| MN.3.06 | H | 2-4 cm | SRR8166127 | 190094 | SRR8166172 | 14882 |
| MN.3.07 | H | 4-6 cm | SRR8166130 | 243958 | SRR8166169 | 17858 |
| MN.3.08 | H | 4-6 cm | SRR8166129 | 74995 | SRR8166170 | 8868 |
| MN.3.09 | H | 4-6 cm | SRR8166132 | 69399 | SRR8166175 | 967 |
| MN.3.10 | H | 6-8 cm | SRR8166131 | 202940 | SRR8166176 | 5613 |
| MN.3.11 | H | 6-8 cm | SRR8166134 | 103760 | SRR8166173 | 7173 |
| MN.3.12 | H | 6-8 cm | SRR8166133 | 222575 | SRR8166174 | 12185 |
| MN.3.13 | H | 8-10 cm | SRR8166124 | 51414 | SRR8166167 | 7688 |
| MN.3.14 | H | 8-10 cm | SRR8166125 | 129387 | SRR8166168 | 5553 |
| MN.3.15 | H | 8-10 cm | SRR8166091 | 118588 | SRR8166190 | 16942 |
| MN.3.16 | H | 10-20 cm | SRR8166106 | 48371 | SRR8166189 | 1841 |
| MN.3.17 | H | 10-20 cm | SRR8166103 | 43541 | SRR8166192 | 1452 |
| MN.3.18 | H | 10-20 cm | SRR8166104 | 113745 | SRR8166191 | 7548 |

[a] Not used because sequence reads were too small (<500 reads)

**Table S3.** Number of sequences obtained by the shotgun metagenome sequencing analyses (GenBank BioProject number PRJNA504043). Short read archive (SRA) accession numbers are also shown for each sample.

| Sample ID | Site | Depth | Metagenome | |
|---|---|---|---|---|
| | | | Accession | No. of Seqs |
| MN.1.01 | M | 0-2 cm | SRX10172081 | 33180362 |
| MN.1.02 | M | 0-2 cm | SRX10172082 | 26353676 |
| MN.1.03 | M | 0-2 cm | SRX10172083 | 34685843 |
| MN.2.01 | I | 0-2 cm | SRX10172084 | 31763677 |
| MN.2.03 | I | 0-2 cm | SRX10172085 | 38328587 |
| MN.3.01 | H | 0-2 cm | SRX10172086 | 39310405 |
| MN.3.02 | H | 0-2 cm | SRX10172087 | 36273765 |
| MN.3.03 | H | 0-2 cm | SRX10172088 | 39798170 |

**Table S4.** Differences in the abundances of archaeal/bacterial 16S rRNA gene and fungal ITS region between the surface soil (0-2 cm) and soils at 8-10 cm or 10-20 cm. Mean +/- SD is shown (n=3). Statistical significance was examined by using Kruskal-Wallis test.

| Parameter | Comparison | Site | | | *p* value[a] |
|---|---|---|---|---|---|
| | | M | I | H | |
| 16S rRNA gene (log copies/g soil) | 0-2 cm vs. 8-10 cm | 2.36 +/- 0.52 | 1.15 +/- 0.61 | 0.78 +/- 0.29 | <0.05 |
| | 0-2 cm vs. 10-20 cm | 2.72 +/- 0.65 | 2.75 +/- 0.65 | 2.90 +/- 0.33 | n.s. |
| Fungal ITS (log copies/g soil) | 0-2 cm vs. 8-10 cm | 5.12 +/- 2.00 | 0.91 +/- 0.39 | 0.98 +/- 0.70 | <0.05 |
| | 0-2 cm vs. 10-20 cm | 4.91 +/- 1.01 | 3.29 +/- 0.01 | 3.72 +/- 1.64 | n.s. |

[a] n.s., not significant

**Table S5.** Differences in the numbers of observed OTUs and the Shannon index values for archaeal/bacterial 16S rRNA gene and fungal ITS region between the surface soil (0-2 cm) and soils at 8-10 cm or 10-20 cm. Mean +/- SD is shown (n=3). Statistical significance was examined by using Kruskal-Wallis test.

| Parameter | Comparison | Site | | | *p* value[a] |
|---|---|---|---|---|---|
| | | M | I | H | |
| No. of obs. OTUs for 16S rRNA gene | 0-2 cm vs. 8-10 cm | 1134 +/- 145 | 489 +/- 288 | 307 +/- 220 | <0.05 |
| | 0-2 cm vs. 10-20 cm | 1495 +/- 472 | 1156 +/- 496 | 1755 +/- 618 | n.s. |
| No. of obs. OTUs for ITS region | 0-2 cm vs. 8-10 cm | 2 +/- 7 | 27 +/- 21 | 4 +/- 18 | n.s. |
| | 0-2 cm vs. 10-20 cm | 14 +/- 20 | 46 +/- 21 | 23 +/- 22 | n.s. |
| Shannon index for 16S rRNA gene | 0-2 cm vs. 8-10 cm | 0.92 +/- 0.20 | 0.34 +/- 0.28 | 0.23 +/- 0.14 | <0.05 |
| | 0-2 cm vs. 10-20 cm | 1.21 +/- 0.28 | 1.45 +/- 0.52 | 1.47 +/- 0.52 | n.s. |
| Shannon index for ITS region | 0-2 cm vs. 8-10 cm | -0.25 +/- 0.12 | 0.56 +/- 0.17 | 0.00 +/- 1.17 | n.s. |
| | 0-2 cm vs. 10-20 cm | -0.44 +/- 0.74 | 1.94 +/- 0.88 | 0.86 +/- 1.35 | n.s. |

[a] n.s., not significant

**Table S6.** Taxa that increased or decreased their relative abundances after the invasion of earthworms identified by Spearman's rank correlation analysis ($p$ <0.01)

| Taxonomic Level | | Taxon | Spearman's ρ |
|---|---|---|---|
| Archaea | Phylum | *Crenarchaeota* | -0.61 |
| | Class | *Thaumarchaeota* | -0.62 |
| | Order | *Nitrososphaerales* | -0.65 |
| | Family | *Nitrososphaeraceae* | -0.65 |
| | Genus | *Nitrososphaera* | -0.60 |
| Bacteria | Phylum | *Actinobacteria* | 0.66 |
| | | *Nitrospirae* | -0.51 |
| | | *Chloroflexi* | -0.50 |
| | Class | *Anaerolineae* | -0.68 |
| | | *Chloracidobacteria* | -0.63 |
| | | *Actinobacteria* | 0.61 |
| | | *Betaproteobacteria* | 0.56 |
| | | *Thermoleophilia* | 0.54 |
| | | *Nitrospira* | -0.51 |
| | Order | *Actinomycetales* | 0.62 |
| | | *Nitrospirales* | -0.51 |
| | Family | *Mycobacteriaceae* | 0.56 |
| | | *Nitrospiraceae* | -0.51 |
| | Genus | *Mycobacterium* | 0.56 |
| | | *Nitrospira* | -0.50 |
| Fungi | Phylum | *Basidiomycota* | 0.47 |
| | | *Ascomycota* | -0.40 |
| | Class | *Leotiomycetes* | -0.59 |
| | Order | *Helotiales* | -0.57 |

**Table S7.** Correlations between the levels of earthworm invasion and the relative abundance of the N cycle-associated genes quantified by the shotgun metagenomics. Spearman's rank-sum test was used for the analysis.

| Gene Functions | Spearman's ρ | *p* value |
|---|---|---|
| Nitrification | | |
| Ammonia monooxygenase (*amo*) | -0.82 | <0.05 |
| Archaeal ammonia monooxygenase (*amo*) | -0.88 | <0.01 |
| Hydroxylamine dehydrogenase (*hao*) | -0.38 | 0.355 |
| Nitrite oxidoreductase (*nxr*) | -0.57 | 0.143 |
| Nitrate reduction | | |
| Nitrate reductase (*nap*, *nar*) | 0.76 | <0.05 |
| Denitrification | | |
| Nitrite reductase (*nir*) | 0.57 | 0.143 |
| Nitric oxide reductase (*nor*) | 0.82 | <0.05 |
| Nitrous oxide reductase (*nos*) | 0.82 | <0.05 |
| Fungal nitrite reductase (fungal *nirK*) | 0.69 | 0.057 |
| Cytochrome P450 nitric oxide reductase | 0.06 | 0.882 |
| DNRA | | |
| Nitrite reductase (*nrf*) | -0.06 | 0.882 |
| Anammox | | |
| Hydroxylamine dehydrogenase (*hdh*) | -0.31 | 0.447 |
| N fixation | | |
| Nitrogenase (*nif*) | 0.88 | <0.01 |
| Assimilation | | |
| Assimilatory nitrate reductase | 0.70 | 0.057 |
| Asparagine synthase | -0.69 | 0.057 |
| Glutamine synthetase | 0.44 | 0.274 |
| Glutamate dehydrogenase | 0.76 | <0.05 |
| Other | | |
| Urease | 0.06 | 0.882 |
| Nitroalkane oxidase | 0.25 | 0.547 |
| Nitronate monoxygenase | 0.19 | 0.654 |
| Glutaminase | 0.38 | 0.356 |

**Table S8.** Fungal trophic modes and functional guilds that increased or decreased their relative abundances after the invasion of earthworms identified by Spearman's rank correlation analysis ($p$ <0.01).

| Fungal trophic modes and functional guilds | Spearman's ρ |
|---|---|
| Trophic mode | |
| Pathotroph | -0.63 |
| Saprotroph | -0.46 |
| Symbiotroph | 0.46 |
| Functional guild | |
| Plant_Pathogen / Undefined_Parasite / Undefined_Saprotroph | 0.56 |
| Plant_Pathogen | -0.54 |
| Ectomycorrhizal | 0.53 |
| Dung_Saprotroph / Ectomycorrhizal / Soil_Saprotroph / Wood_Saprotroph | 0.52 |
| Endomycorrhizal / Plant_Pathogen / Undefined_Saprotroph | 0.50 |
| Dung_Saprotroph / Ectomycorrhizal / Litter_Saprotroph / Undefined_Saprotroph | 0.44 |
| Dung_Saprotroph / Soil_Saprotroph / Undefined_Saprotroph | -0.43 |
| Undefined_Saprotroph | -0.41 |

**Supplementary Figures**

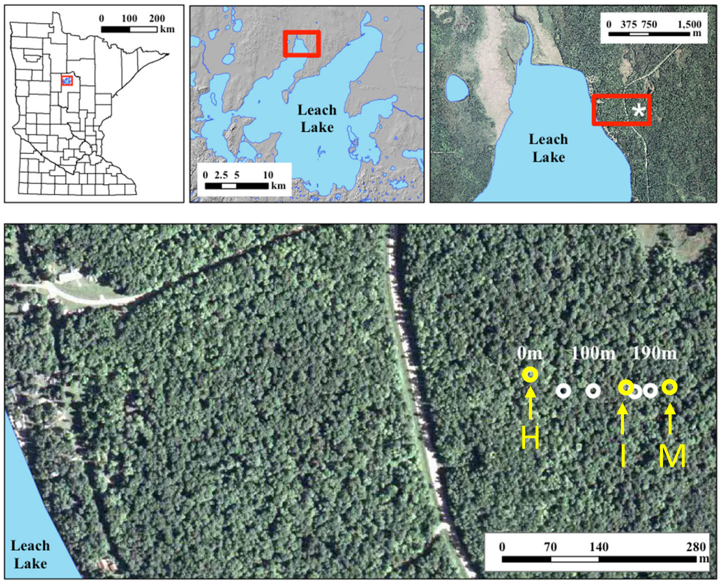


**Figure S1.** Maps of study area near Leech Lake, Minnesota and the sampling sites along the earthworm invasion chronosequence (adapted from [4]). Legend: H, the highly invaded site; I, the intermediately invaded site; M, the minimally invaded site.

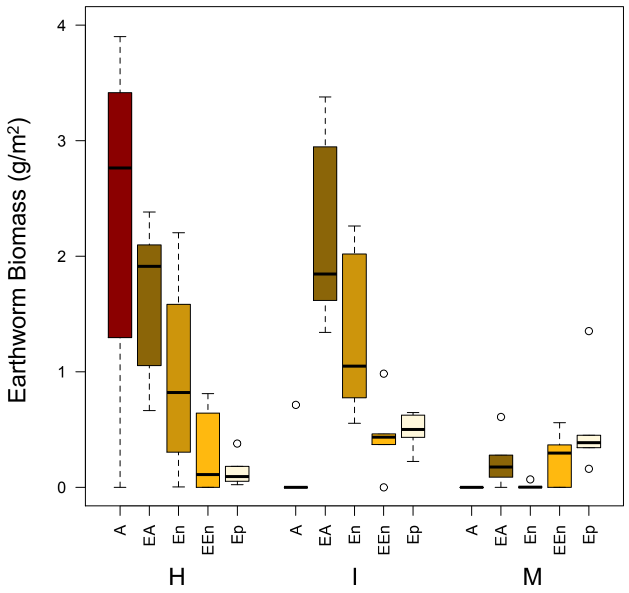


**Figure S2.** Biomass of earthworms of different types. Legend: A, anecic earthworms (adult *Lumbricus terrestris*); EA, epi-anecic earthworms (juvenile *Lumbricus terrestris*); En, endogeic earthworms (*Aporrectodea* spp. and *Octolasion* spp.); EEn, epi-endogeic earthworms (*Lumbricus rubellus*); Ep, epigeic earthworms (*Dendrobaena* spp.); H, the highly invaded site; I, the intermediately invaded site; M, the minimally invaded site.

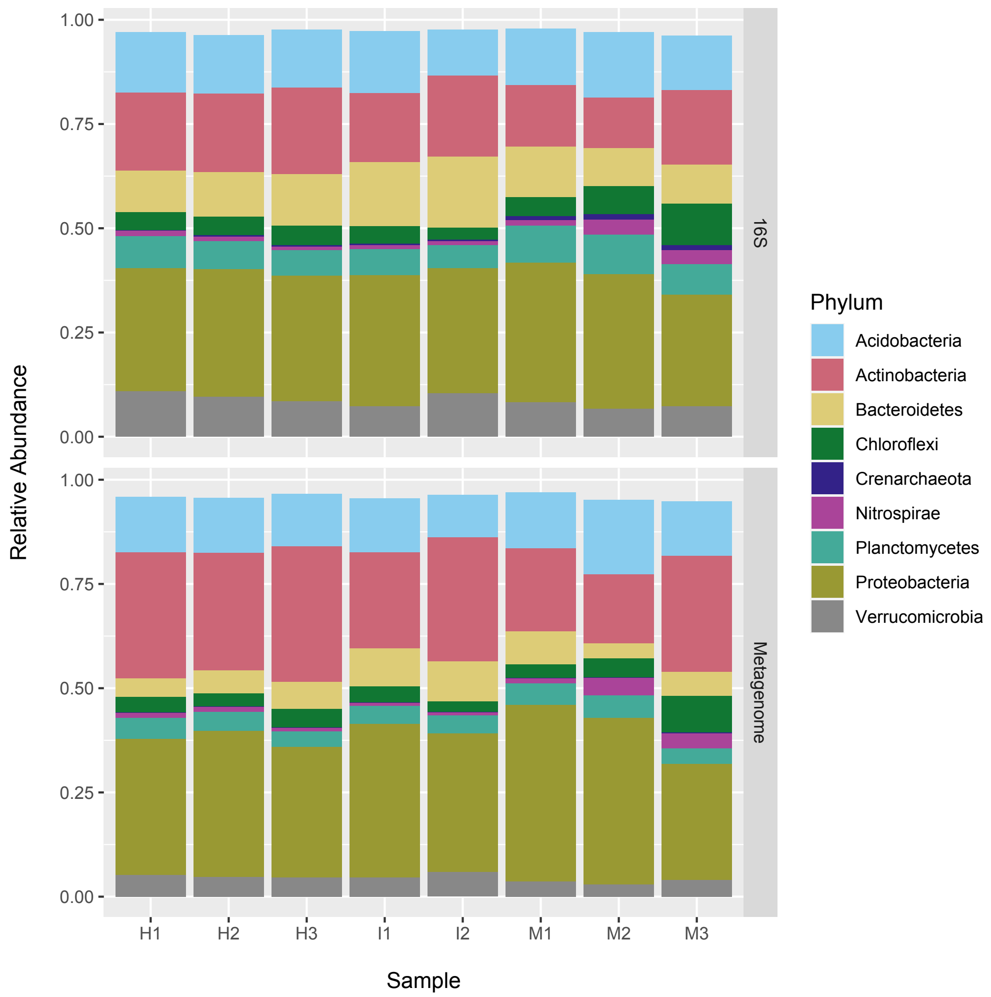


**Figure S3.** Comparison of the soil microbiomes (taxonomic compositions at a phylum level) analyzed by the 16S rRNA gene amplicon sequencing and the shotgun metagenomics. Soil samples collected from the surface (0-2 cm depth) were used. Legend: H1, the highly invaded site replicate 1; H2, the highly invaded site replicate 2; H3, the highly invaded site replicate 3; I1, the intermediately invaded site replicate 1; I2, the intermediately invaded site replicate 2; M1, the minimally invaded site replicate 1; M2, the minimally invaded site replicate 2; M3, the minimally invaded site replicate 3.

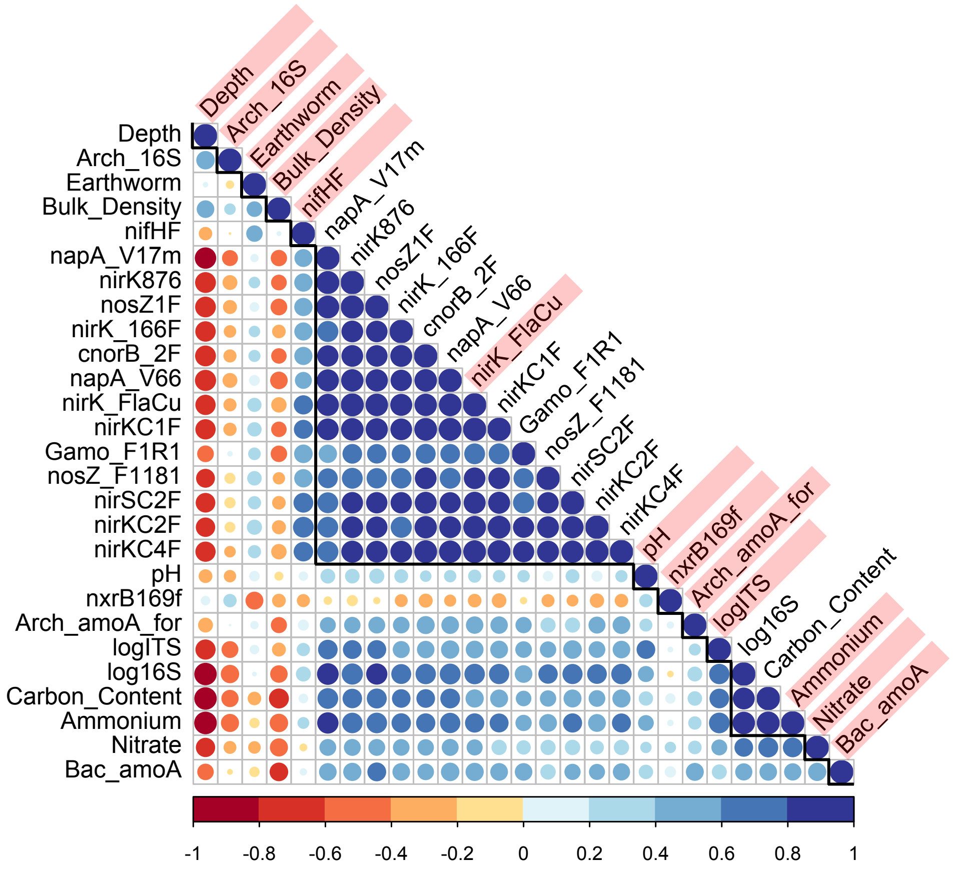


**Figure S4.** Correlations between the environmental variables measured in this study. Positive and negative correlations are shown in blue and red, respectively. The color intensity and the size of the circle are proportional to Spearman's ρ value. Environmental variables shaded with pink were selected and used for the CAP analysis.
